# Liquid-like and rigid-body motions in molecular-dynamics simulations of a crystalline protein

**DOI:** 10.1101/811083

**Authors:** David C. Wych, James S. Fraser, David L. Mobley, Michael E. Wall

## Abstract

To gain insight into crystalline protein dynamics, we performed molecular-dynamics (MD) simulations of a periodic 2×2×2 supercell of staphylococcal nuclease. We used the resulting MD trajectories to simulate X-ray diffraction and to study collective motions. The agreement of simulated X-ray diffraction with the data is comparable to previous MD simulation studies. We studied collective motions by analyzing statistically the covariance of alpha-carbon position displacements. The covariance decreases exponentially with the distance between atoms, which is consistent with a liquid-like motions (LLM) model, in which the protein behaves like a soft material. To gain finer insight into the collective motions, we examined the covariance behavior within a protein molecule (intra-protein) and between different protein molecules (inter-protein). The inter-protein atom pairs, which dominate the overall statistics, exhibit LLM behavior; however, the intra-protein pairs exhibit behavior that is consistent with a superposition of LLM and rigid-body motions (RBM). Our results indicate that LLM behavior of global dynamics is present in MD simulations of a protein crystal. They also show that RBM behavior is detectable in the simulations but that it is subsumed by the LLM behavior. Finally the results provide clues about how correlated motions of atom pairs both within and across proteins might manifest in diffraction data. Overall our findings increase our understanding of the connection between molecular motions and diffraction data, and therefore advance efforts to extract information about functionally important motions from crystallography experiments.

## Introduction

Macromolecular crystals consist of many copies of a large molecule (or molecules) packed into a lattice of repeating units. Crystals are often illustrated using identical repeating units; however, in real crystals, each copy of a molecule can adopt a somewhat different structure as long as overall order is maintained. Although structural variations occur in small-molecule crystals^1^, the problem of conformational heterogeneity is especially important for macromolecular crystals, which have many more degrees of freedom and high solvent content.^2 3^ Understanding the structural variations in crystals can potentially be used for increasing the accuracy of crystal structure models^4^ and for developing crystallography as a tool for characterizing conformational heterogeneity and dynamics in structural biology.^5^

In X-ray crystallography (and similarly for neutron and electron crystallography), the details of the molecules’ conformations and their packing in the crystal lattice leave signatures in the diffraction pattern. Most of our current understanding of conformational variation in crystals comes from analysis of Bragg reflections, which are the sharp peaks in the diffraction pattern. The Bragg peaks are traditionally modeled using an average picture of the repeating unit, or unit cell, which contains the mean electron density over all unit cells illuminated by the beam. The most common model multiplies each atomic form factor by an atomic displacement parameter (also known as a Debye-Waller factor, B-factor, or thermal factor) corresponding to a 3D Gaussian distribution of each atom’s displacements.

Modern Bragg analysis methods are producing richer descriptions of conformational variations. For example, anisotropic displacements of groups of atoms can be modeled with a small number of parameters using the Translation Libration Screw (TLS) model,^6^ which can be used as a supplement or substitute to refining individual atomic displacement parameters.^7^ More elaborate models of conformational heterogeneity can also be employed,^8^ including incorporating local alternative conformations in multiconformer models^9 10^ or generating multiple models using hybrid molecular dynamics ensemble refinement that collectively satisfy the data.^10^ These models can imply distinct collective motions that would leave signatures in the spatial correlations of electron density variations. Therefore, even if more elaborate model of motion improve agreement with the Bragg data, they would need to be validated to determine whether they describe what is actually happening in the crystal.^11^ Indeed, distinct TLS refinements with different implied collective motions can yield equivalent agreement with the Bragg data.^12^

The degeneracy of models with respect to the mean electron density fortunately does not extend to spatial correlations in electron density variations. Models with different correlations lead to distinct patterns of diffuse scattering that appear as intensity underneath and between the Bragg peaks in diffraction images. Recent years have seen a renaissance in methods for processing and modeling diffuse scattering.^3 2 13^ However, studies of diffuse scattering have differed in their conclusions, both about the degree to which the signal can be explained by various models of correlated motion, and about the importance of different types of motion in determining the conformational variations in protein crystals.^14 15 16 2 17^ Some studies find evidence for liquid-like motions (LLM), which models the crystal contents as a soft material with correlated displacements on a characteristic length scale.^18^ Others find evidence for rigid-body motions (RBM),^15^ which could, in principle, be modeled by descriptions such as TLS models.^12^ Ensemble models, in which a limited set of representative structures are selected from a full conformational distribution,^19^ also have been studied, and found lacking.^14 15^ A potential problem with ensemble models is that they exhibit exaggerated correlations due to the absence of finer-scale structure variations.^20^ More sophisticated models might be able to better capture the correlated motions within the crystal.^21^ Considering the true crystalline context by including neighboring unit cells in the calculations can increase agreement of simple models like LLM.^14^

Whereas each of the above mentioned models of diffuse scattering explicitly represents only a subset of the possible structural variations, molecular-dynamics (MD) simulations of crystalline proteins provide an all-atom picture of a wide variety of available motions.^22 23 24 25^ Early MD simulations of diffuse scattering were hindered by the short duration (< 1 ns) which resulted in limited sampling of the conformational ensemble, and calculations that were not reproducible across different runs.^26^ Agreement between experimental and predicted diffuse scattering of crystalline staphylococcal nuclease increased as the simulation duration increased to 10 ns^27^ and then even more as it was increased to 1 microsecond.^25^ A recent simulation of staphylococcal nuclease extended the simulation volume from a single periodic unit cell to a 2×2×2 supercell, leading to further increases in the agreement with the experimental data.^28^ This result parallels the emerging theme from the LLM work,^14^ which highlighted the importance of considering more than just a single unit cell to generate strong agreement with experimental data.^29^ MD provides a full description of all atoms in the system, but has a high computational cost relative to the simpler models, such as LLM and RBM. Because simpler models can potentially be incorporated into joint X-ray Bragg and diffuse scattering model refinement, it is interesting to investigate the degree to which the motions described by simpler models, such as the LLM and RBM models, appear in the increasingly accurate MD simulations that are now achievable. It is worth noting that while the LLM and RBM models include parameters explicitly fit to match diffuse scattering patterns, MD models utilize no such fit, making this comparison particularly interesting.

Here we assess the degree to which LLM and RBM behaviors are present in MD simulations of crystalline staphylococcal nuclease. We performed new MD simulations of crystalline staphyloccocal nuclease and confirmed their agreement with diffuse scattering data is similar to previous studies. We then computed covariance matrices of atom displacements from the resulting MD trajectories, and analyzed the dependence of the covariance on the average separation between atoms which are covarying. The covariance behavior is well fit by an exponential decrease with distance, supporting a LLM model. However, when the analysis is restricted to atom pairs that lie within the same protein, the behavior is consistent with a combination of LLM and RBM. Comparison of results obtained using AMBER vs CHARMM force fields reveal some differences in the simulations and covariance behavior. Our results indicate that LLM behavior of global dynamics emerges from MD simulations of a protein crystal. They also show that RBM behavior is detectable in the simulations but that it is subsumed by the LLM behavior when adding atom pairs that cross protein boundaries. Finally they provide clues about how correlated motions of atom pairs both within and across proteins might manifest in the diffuse scattering data. Overall our findings increase our understanding of the connection between molecular motions and diffuse scattering data, and therefore advance efforts to extract information about functionally important motions from crystallography experiments.

## Results

### Molecular dynamics simulations

In this study we sought to investigate the connection between MD simulations and other, less detailed models of crystalline dynamics, like the LLM model. For the MD simulations, we used a model of crystalline staphylococcal nuclease, consisting of a periodic box consisting of 2×2×2 unit cells (Fig. 1; Methods). Solution state simulations commonly are performed using a NPT ensemble; however, as we wished to compare our simulations to diffraction data, we used a NVT ensemble, maintaining the consistency of the system with the experimentally determined Bragg lattice during the course of the simulation. To determine the sensitivity of the results to the force field, we used both AMBER 14SB and CHARMM 27 force fields in GROMACS.

**Figure 1.**
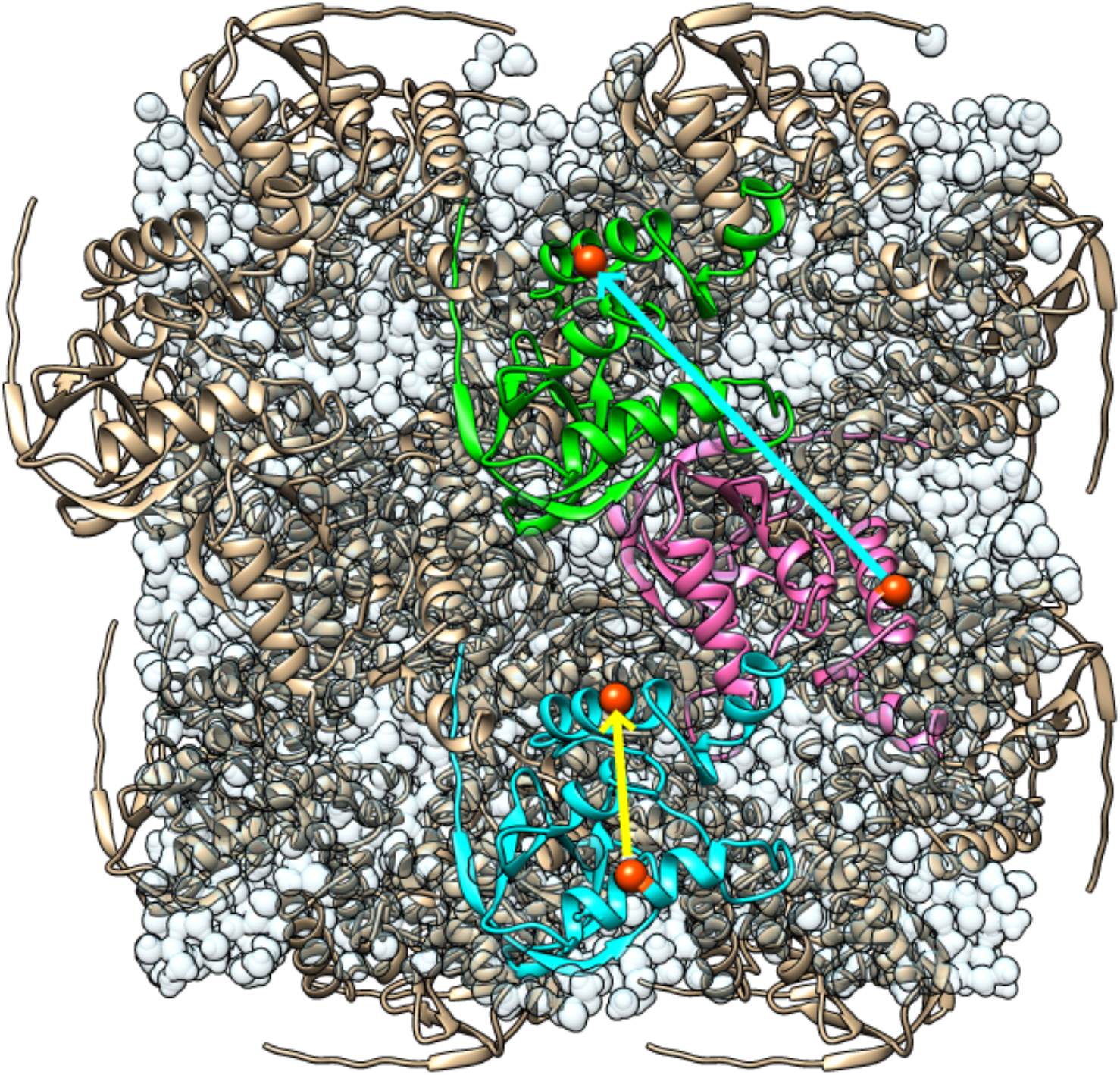
Illustration of the model used for molecular dynamics simulations. There are 32 copies of the protein rendered using ribbons in a periodic box of 2×2×2 unit cells. The yellow arrow indicates an atom pair that lies entirely within the blue protein, pointing from residue 61 to residue 131 (corresponding to the “within” or “intra-protein” analysis). The cyan arrow indicates an atom pair that spans across proteins, pointing from residue 131 in the magenta to residue 128 in the green protein (corresponding to the “across” or “inter-protein” analysis). Water molecules are rendered in light, transparent blue, giving the appearance of connected droplets. The image was created using UCSF *Chimera*.^30^

Consistent with previous studies,^28 25^ which used similar methods to the present study, after initial solvation and minimization, during the first equilibration step, the pressure of the systems was large and negative: −1439 +/− 39 bar for the AMBER simulation, and −1795 +/− 252 bar for the CHARMM simulation (as reported by *gmx energy*). To bring the system to atmospheric pressure, additional water molecules were added iteratively, with intervening rounds of NVT equilibration, until the mean system pressure was in the range of −100 to 100 bar (Methods). The number of water molecules added was roughly the same for both systems (17,557 waters for AMBER and 17,138 waters for CHARMM).

After the iterative solvation steps, unrestrained MD simulations were carried out for 600 ns. The potential energy drifted during the first ~100 ns the simulation, after restraints were released, and remained relatively stable thereafter. To ensure that the drift did not influence the results of our analysis of the trajectory, the initial 200 ns section was ignored, and only the last 400 ns section was analyzed. The B-factors predicted by the atomic fluctuations (not shown) were consistent with previous studies of the same system ^28^, and with the experimentally determined B factors.

### Simulated diffuse intensity

To determine the agreement of the simulations with diffuse scattering data, we computed diffuse intensities from MD trajectories and computed the Pearson correlation coefficient between the simulated and experimental intensities (Methods). Simulated diffuse diffraction images computed from the MD simulations and experimental data are compared in Fig. 2. The Pearson correlation between the simulated and experimental total 3D diffuse intensities was 0.9 or higher for all simulations, similar to that found in a previous study utilizing the same approach.^28^ The correlation of the anisotropic component of the intensity, computed by subtracting an interpolated radial average,^28^ was 0.58 (AMBER) and 0.63 (CHARMM) for the anisotropic intensity, which is lower than the value of 0.68 previously reported in Ref. 28. We re-analyzed the data in Ref. 28 and obtained the same correlation of 0.68, confirming that the difference is attributed to the simulations rather than the analysis workflow.

**Figure 2.**
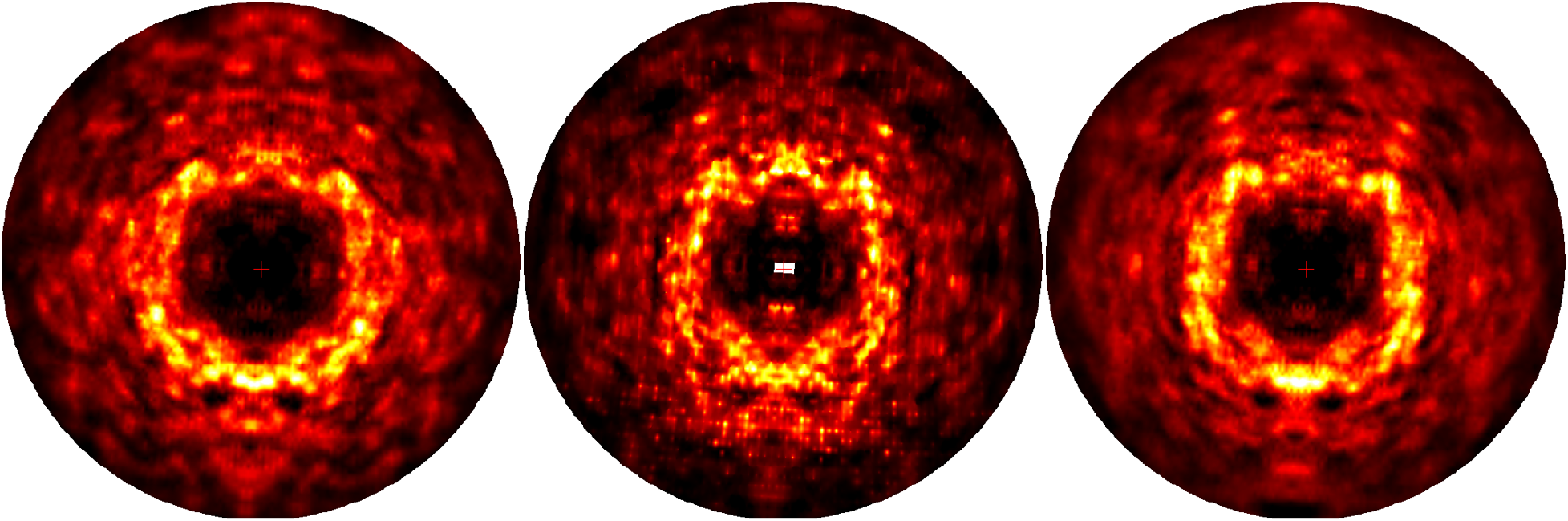
Simulated diffraction images derived from MD simulations and 3D experimental diffuse data. 2×2×2 sampling. The images are truncated at 1.8 Å. To make the anisotropic features more visible, the 3D diffuse intensities have had the minimum intensity value subtracted in constant resolution shells prior to generating these images. *Left.* Image derived from AMBER force-field simulation. *Center.* Image derived from experimental data. *Right.* Image derived from CHARMM force-field simulation. The diffuse intensity for MD simulations was accumulated incoherently across 100 ns sections of the trajectory in the time range 200 - 600 ns after relaxing restraints. Both the AMBER and CHARMM MD simulations show similar diffuse features to those in the experimentally derived images, e.g., the strong intensity in the ring and cloudy features at higher resolution. The images were displayed using the heat map mode of ADXV.^31^

To gain finer-grained insight into the agreement between the simulated and experimental diffuse intensities, we analyzed the trajectory in 100 ns sections (Supplementary Fig. S1). The correlation computed using each section is between 0.53 and 0.58 which is lower than the 0.58 and 0.63 values obtained using the diffuse intensity accumulated for 400 ns. The cumulative agreement also is sensitive to whether the diffuse intensity is accumulated coherently or incoherently across the individual 100 ns sections (see Methods for definition of coherent vs incoherent accumulation): when accumulated coherently, the correlation is 0.54 (AMBER) 0.58 (CHARMM) which is lower than the values 0.58 (AMBER) and 0.62 (CHARMM) when accumulated incoherently.

### Liquid-like motions behavior of all Cα atom pairs

To assess whether the MD simulations produced behavior that is consistent with the LLM model, we computed atom displacement covariance matrices (Methods). The covariance matrices are needed because the key assumption of the LLM model is that the covariance matrix elements connecting any two atoms *i* and *j* in the crystal are simply proportional to 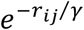, where *r*_*ij*_ is the distance between the atoms, and *γ* is the characteristic length scale of the correlations.^18^ To make the calculations and analysis manageable computationally and to obtain a coarse-grained picture of the covariance, we restricted the analysis to Cα atoms.

The full covariance matrix of Cα displacements for our system is a 14,304 x 14,304 square matrix with each row or column corresponding to one of three cartesian coordinates of each of 149 Cα atoms in each of 32 proteins in the model. To perform our analyses, we replaced the 3×3 submatrix for each atom pair with its trace, leaving a 4,768 x 4,768 square symmetric matrix. In this form, the diagonal elements correspond to the mean squared deviation (MSD) for each atom, and the off-diagonal elements correspond to the trace of the covariance of the displacements of each atom pair. The full covariance matrix contains regions of both positive and negative covariance, with the strongest positive values being in blocks about the diagonal (Supplementary Fig. S2), corresponding to atom pairs that fall within a protein (as connected by the yellow line in Fig. 1).

To determine the dependence of the matrix elements coupling atoms *i* and *j* on the distance between the atoms *r*_*ij*_, we computed the distance between each of the atom pairs and divided the distance range into 50 even bins. We then calculated the mean and standard error of the covariance within each of the bins (Methods). For each simulation, at the lowest distance there is a single bin with high covariance compared to the other bins: in the AMBER case, a point at 1.67 Å with covariance 15.5 Å^2^ +/− 1.84 Å^2^ as it is out of range in y; in the CHARMM case, a point at 1.47 Å with covariance 21.8 Å^2^ +/− 3.06 Å^2^ is not shown, as it is out of range in y. Beyond 5 Å (Fig. 3) the covariance is much lower (20-fold lower than in the nearest bin below) and shows a more gradual decrease with distance, falling from a value of 0.30 Å^2^ at 5 Å, crossing below zero beyond about 40 Å to a minimum value of either −0.02 (AMBER) or −0.03 (CHARMM) Å^2^ beyond about 50 Å, and rising to a value closer to zero beyond about 80 Å (AMBER) or 60 Å (CHARMM).

**Figure 3.**
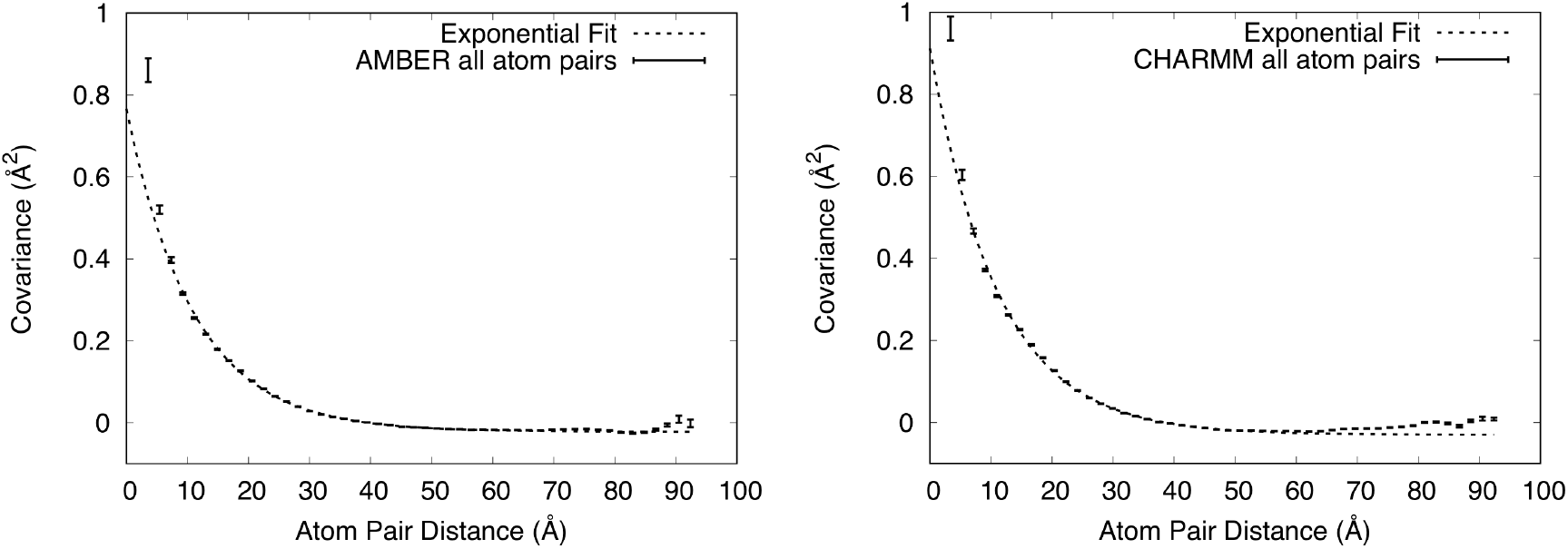
Dependence of covariance on distance for all Cα atom pairs. Mean values of covariance +/− standard errors are shown using vertically oriented error bars. Exponential fits to the values in the 5-55.5 Å range are shown using dashed lines. The upper y-range is truncated at 1 Å^2^, showing the full range of the exponential fit but excluding one high covariance value at very short distance from each panel. *Left.* AMBER force field. The exponential fit has a decay length of 11.0 Å. *Right.* CHARMM force field. The exponential fit has a decay length of 11.1 Å.

To assess whether these plots display exponential decay behavior, the values of the covariance *C*(*r*) were fit to the function *C*(*r*) = *ae*^−*r*/*γ*^ + *b*, where *r* is the distance between atoms, in the range *r* between 5 Å and 55.5 Å. (We found the exponential fit was poor without adding the constant *b*, so we added it.) For the AMBER simulation, the fit yielded *a* = 0.79 ± 0.01 Å^2^, γ = 11.0 ± 0.1 Å, and *b* = −0.022 ± 0.001 Å^2^. For the CHARMM simulation, the fit yielded *a* = 0.94 ± 0.02 Å^2^, γ = 11.1 ± 0.2 Å, and *b* = −0.029 ± 0.001 Å^2^. The constant offset at long distances is consistent with an earlier simulation,^27^ which postulated that it might be an artifact of translational alignment of the MD trajectory snapshots – a necessary step before computing the covariance matrix (Methods). Fig. 3 shows that the fit overlaps the computed covariances in the region below 60 Å, and therefore displays exponential decay behavior, which is consistent with the assumption of the LLM.

### Combination of liquid-like and rigid-body motions behaviors within proteins

As noted in the previous section, the strongest positive values of the covariance matrix were in blocks along the diagonal (Supplementary Fig. S2). These blocks correspond to Cα atom pairs that lie within the same protein (yellow line in Fig. 1), which we refer to as “intra-protein” or “within-protein” atom pairs. The rest of the covariance matrix corresponds to atom pairs that cross protein boundaries, which we refer to as “inter-protein” or “across-protein” atom pairs (cyan line in Fig. 1). (The subsets of intra-protein and inter-protein Cα atom pairs are complementary with respect to the set of all Cα atom pairs.) We wondered whether the LLM behavior observed for all Cα atom pairs also would apply individually to these subsets. We therefore computed the covariance vs distance for each. We also wished to focus our attention on the more stable region of the protein and therefore eliminated residues at the extreme N- and C-terminal Cα atoms from our calculations (Methods).

The shape of the curve for inter-protein atom pairs (Fig. 4) is similar to that for all Cα atom pairs (Fig. 3). In the case of the AMBER simulation, a point at 3.3 Å with covariance −1.77 Å^2^ +/− 1.23 Å^2^ is not shown as it is out of range in y. For AMBER the best-fit exponential decay has *a* = 0.42 ± 0.02 Å^2^, γ = 14.3 ± 0.4 Å, and *b* = 0.025 ± 0.001 Å^2^; for CHARMM the best-fit has *a* = 0.55 ± 0.02 Å^2^, γ = 13.4 ± 0.3 Å, and *b* = 0.032 ± 0.001 Å^2^. The values at short distance are smaller in the inter-protein analysis than in the analysis of all atom pairs. In the case of the AMBER simulation, the exponential fit deviates from the covariance values in the region below 10 Å (Fig. 4). The values of *a* are smaller than for all atom pairs, supporting the observation that the values at short distance are smaller. The values of γ are larger than for all atom pairs, indicating that correlations extend to a longer length scale. The values of *b* are similar to the values for all atom pairs.

**Figure 4.**
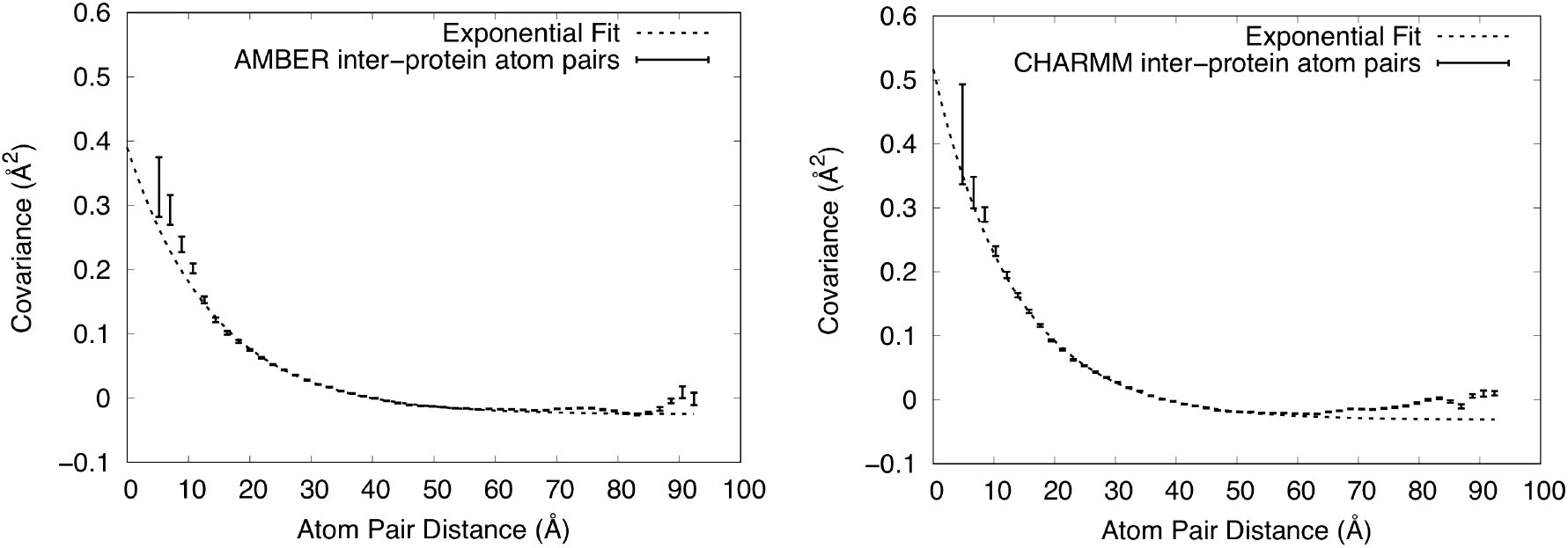
Dependence of covariance on distance for just inter-protein Cα atom pairs. Mean values of covariance +/− standard errors are shown using vertically oriented error bars. Exponential fits to the values in the 5-55.5 Å range are shown using dashed lines. The upper y-range is truncated at 0.6 Å^2^. *Left*. AMBER force field. The shortest distance point is not shown as it is out of range. The exponential fit has a decay length of 14.3 ± 0.4 Å. *Right*. CHARMM force field. The exponential fit has a decay length of 13.4 ± 0.3 Å. The exponential fits are fairly good, but not as good as in Figs. 3.

In contrast with the inter-protein atom pairs, the shape of the curve for atom pairs within proteins (Fig. 5) is very different (note that the x-axis only extends to ~42.5 Å, while retaining the number of bins at 50). In the AMBER case, a MD point at 2.90 Å with covariance 1.64 Å^2^ +/− 0.38 Å^2^ is not shown as it is out of range in y. In the case of the CHARMM simulation, A MD point at 3.2 Å with covariance 2.85 Å^2^ +/− 0.23 Å^2^ is not shown as it is out of range in y. The values still decrease with increasing distance, but the curvature is lower near 5 Å. Moreover the behavior is almost linear above 20 Å, crossing zero at about 38 Å, and decreasing to about −0.05 Å^2^ at the longest distance.

**Figure 5.**
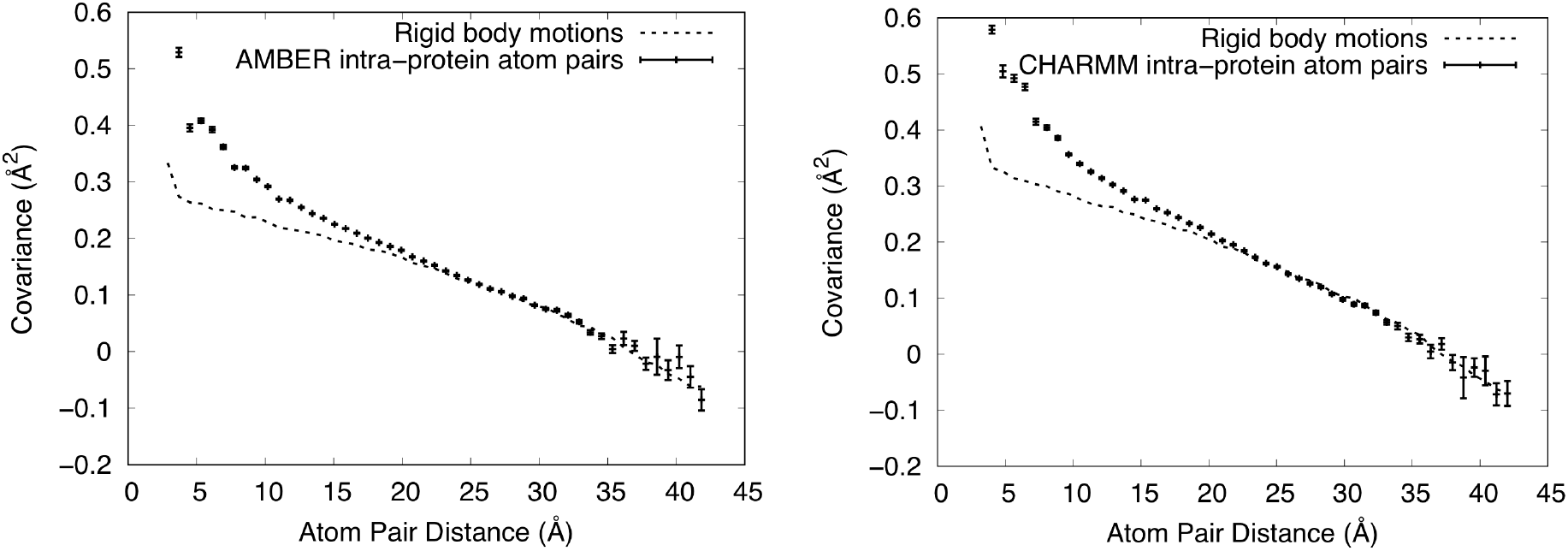
Dependence of covariance on distance for only intra-protein Cα atom pairs. Mean values of covariance +/− standard errors from the MD simulations are shown using vertically oriented error bars. Values computed from RBM models are shown using dashed lines (standard errors are O(10^−5^) Å^2^ and are therefore not shown). The upper y-range is truncated at 0.6 Å^2^, showing the full range of the exponential fit but excluding one high covariance value at the shortest distance from the MD values in each panel. *Left*. AMBER force field. The RBM model has a width (SD) 0.95 degrees for the angular distribution and 0.24 Å for the translational distribution. *Right*. CHARMM force field. The RBM model has a width (SD) 1.05 degrees for the angular distribution and 0.27 Å for the translational distribution. The MD simulations are well modelled using rigid-body translations and rotations for distances greater than 20 Å.

In seeking an explanation for the linear behavior, we reasoned that rigid-body rotations of individual proteins should give rise to a decreasing covariance with distance. For example, during a rotation through the center of mass, nearby atoms on the surface would tend to move together, giving rise to a positive covariance, and atoms at remote locations on the surface would tend to move in opposite directions, giving rise to a negative covariance. We postulated that this might lead to a decreasing covariance with distance that is positive at short distances and becomes negative at long distances. Adding a rigid-body translation after the rotation would lead to a uniform positive covariance within the protein, shifting the curve up and the zero crossing to longer distances.

To test this idea, we generated an ensemble of snapshots displaying varying degrees of RBM. For each snapshot, three Euler angles and a translational shift were drawn from a normal distribution, and rigid coordinate transformations were applied to a single staphylococcal nuclease protein from the MD model. The widths of the distributions were chosen to be on par with the magnitude of motion observed in our MD simulations. The covariance matrix was computed from the snapshots, and the distance dependence was analyzed in the same manner as for the MD trajectories.

We adjusted the standard deviation (SD) of the angular and translational distributions to optimize the visual agreement with the MD simulation in the long-distance part of the curve. In the case of the AMBER simulation, the final SD used for the angular distribution was 0.95 degrees, and the SD of the translational distribution was 0.24 Å. For the CHARMM simulation, the SD of the angular distribution was 1.05 degrees, and the SD of the translational distribution was 0.27 Å. The model tracks MD simulation results in the region above about 20 Å (Fig. 4), indicating that the MD covariance in this long-distance region is consistent with RBM.

To explain the behavior in the region below 20 Å, we subtracted the RBM plots from the corresponding MD simulation plots in Fig. 5, yielding the residual shown in Fig. 6. In the AMBER case, a point at 2.9 Å with covariance 1.31 Å^2^ is not shown as it is out of range in y. In the CHARMM case, a MD point at 3.2 Å with covariance 2.45 Å^2^ is not shown as it is out of range in y. As the residual plots resemble an exponential decay, we again fit them to the function *C*(*r*) = *ae*^−*r*/*γ*^ + *b* in the region *r* > 5 Å. For the AMBER simulation, the fit yielded *a* = 0.37 ± 0.02 Å^2^, γ = 5.7 ± 0.2 Å, *b* = 0 Å^2^ to within error; for CHARMM, the fit yielded *a* = 0.46 ± 0.03 Å^2^, γ = 5.7 ± 0.3 Å, and *b* = −0.002 ± 0.001 Å^2^. The fit confirms that the residual is consistent with an exponential decay, but with a much shorter length scale than for the inter-protein atom pairs.

**Figure 6.**
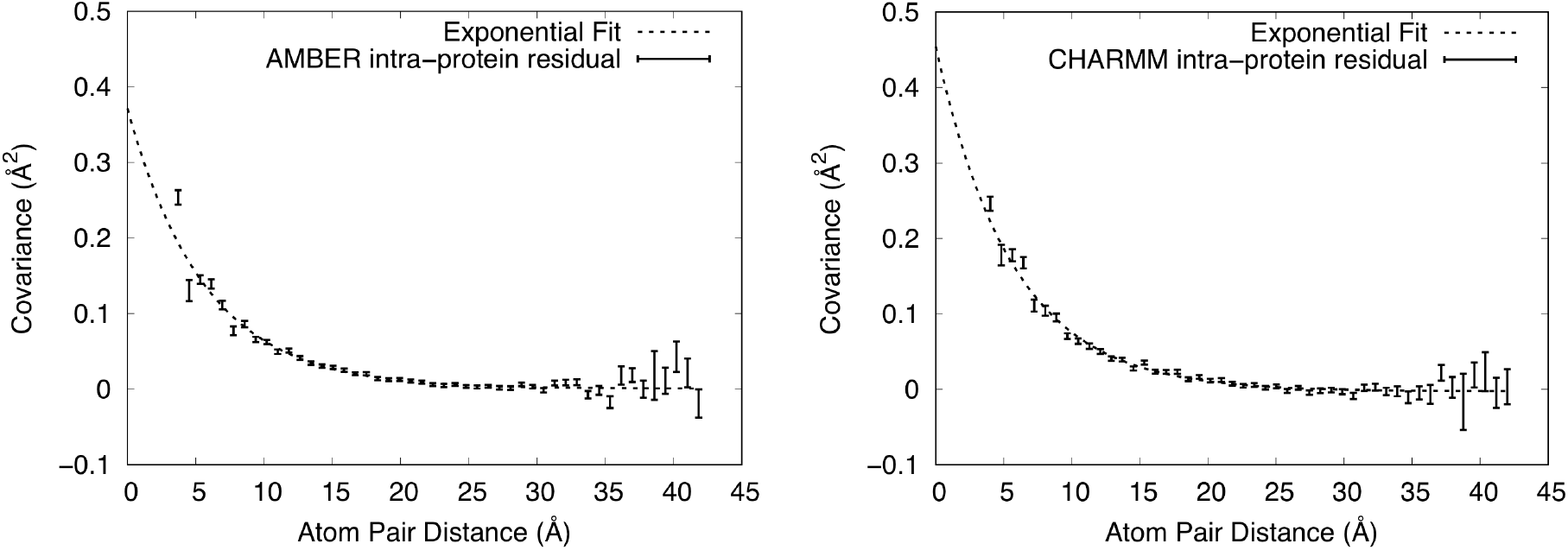
Residual covariance computed from the MD simulations, after subtracting the covariance values of the RBM model (see Fig. 5). Mean values of covariance +/− standard errors from the MD simulations are shown using vertically oriented error bars. Exponential fits to the values in the range above 5 Å are shown using dashed lines. The upper y-range is truncated at 0.5 Å^2^, showing the full range of the exponential fit but excluding one high covariance value at the shortest distance from each panel. *Left*. AMBER force field. *Right*. CHARMM force field. Both residuals are well fit by an exponential with the same decay length of 5.7 Å.

Additional insight into the LLM behavior comes from comparing the values of γ and *a* obtained from the MD analysis with the values obtained by fitting a LLM model to coarsely sampled (one point per Miller index) experimental diffuse scattering data (Methods). The refined LLM model had a Pearson correlation coefficient with the anisotropic data of 0.73, with γ = 6.5 Å and σ = 0.41 Å (the agreement with the total diffuse data, as opposed to the anisotropic data, is poor, as the LLM model does not include solvent). A comparison of simulated diffraction images from the model and data is shown in Supplementary Fig. S4. The value of γ is closer to the value for the intra-protein analysis (5.5 Å) than for the inter-protein (11.5 Å) or all-atom (12 Å) analysis. The value of σ corresponds to a MSD of 3 × (0.41 Å)^2^ = 0.50 Å^2^ which is comparable to the values obtained in the intra-protein analysis (0.42 Å^2^ for AMBER and 0.48 Å^2^ for CHARMM) and to the values for the inter-protein analysis (0.55 Å^2^ for AMBER and 0.66 Å^2^ for CHARMM). This comparison indicates that the coarsely sampled diffuse scattering data exhibit both a length scale of correlations and amplitude of motion that are most consistent with the intra-protein atom pairs in the MD simulation (see Discussion).

## Discussion

Our crystalline MD simulations of staphylococcal nuclease reveal a consistency with the LLM model: the distance dependence of the covariance of Cα displacements follows an exponential decay. This finding indicates that LLM behavior is present in a more realistic, highly detailed, all-atom description of the dynamics. It also provides a rationale for why the LLM model, which uses only a few parameters, can provide a reasonable explanation of diffuse scattering data, which depends on the atomic details of the structure variations.

The atomic details contained in the MD also allow us to examine the covariance behavior within a protein molecule (intra-protein) and between different protein molecules (inter-protein). Consistent with the overall LLM model, the distance dependence of the covariance between inter-protein atom pairs appears exponential, albeit with deviations below 10 Å (Fig. 4). The best-fit values of γ are 14.3 Å for the AMBER simulation and 13.4 Å for the CHARMM simulation, which are somewhat longer than the value of 11 Å for all protein pairs. If the subset of inter-protein atom pairs dominates the statistics of all atom pairs, then it should be most important in determining the LLM behavior that was observed for all atom pairs. Indeed, beyond a distance of 12 Å, the number of inter-protein atom pairs sharply climbs above the number of intra-protein atom pairs (Supplementary Fig. S3), supporting this notion. This finding is consistent with earlier work of Peck et al.^14^ showing that including interactions across molecular boundaries improves agreement with the anisotropic diffuse scattering signal.

In contrast to the inter-protein atom pairs, the covariance behavior for just the intra-protein atom pairs deviates from a LLM model. For these atom pairs, the behavior is described by a mixture of RBM and other contributions. The long-range behavior (above 20 Å) is almost exclusively explained by RBM, the mid-range behavior (5-20 Å) is dominated by RBM, and in the short-range region (below 5 Å) the RBM model accounts for only a minority of the covariance. The observation of RBM here is reminiscent of ssNMR studies of ubiquitin crystals^32, 33^ in which MD simulations were used to explain the crystal dynamics; in that study, 3-5 degree rocking motions were observed via rotational alignment of proteins from the MD trajectory. We note, however, that 3-5 degrees is much larger than the 1 degree SD that explains the covariance behavior in Fig. 5. In addition, the present results are consistent with our previous analysis of rigid-body rotations based on rotational alignment of proteins from the MD trajectory^28^; that analysis found 1-2 degree SDs of Euler angles and indicated that rigid-body rotations account for a minority of the atom displacements in staphylococcal nuclease crystalline MD simulations.

After subtracting the RBM contribution from the intra-protein covariance plot, the residual is well fit by an exponential decay, except below 5 Å. The decay length of the fit is 5.7 Å, which is substantially shorter than the length found for all atom pairs (11 Å) or just the inter-protein atom pairs (13.4 Å and 14.3 Å). As the decay length differs, it is possible that the inter-protein and intra-protein LLM behaviors have different origins in the MD simulations. As the intra-protein LLM behavior only is apparent after the RBM contribution has been subtracted, it is unlikely to involve a substantial RBM component; however, it is possible that the inter-protein LLM behavior includes a component that is due to the coupling of RBM across protein boundaries.^34^

The value γ = 6.5 Å obtained for the LLM fit to staphylococcal nuclease diffuse scattering data is substantially smaller than the 18 Å value obtained by Peck et al.^14^ for LLM models of diffuse scattering from CypA and WrpA. Peck et al.^14^ also noted that their values of γ were larger than previously published values, and that the difference in length scales for LLM models might be attributed to their finer sampling of the data. Our results lend support to this explanation for the discrepancy. In the case of intra-protein atom pairs, we found γ = 5.7 Å, which is comparable to the value γ = 6.5 Å from the LLM fit. In the case of inter-protein atom pairs, we found γ = 14.3 Å (AMBER) or γ = 13.4 Å (CHARMM), which is more comparable to the value γ = 18 Å from the Peck et al.^14^ study. The similarity of these length scales between the fitting and the MD suggests that both types of motions might be present in the protein crystal. Moreover the comparison of the length scales between MD analysis and LLM models suggests that the fine-grained sampling might yield data that emphasize inter-protein motions, and that the coarse-grained sampling might yield data that emphasize intra-protein motions in the fitting. This possibility motivates future work to identify regions of reciprocal space where the intra-protein signal is enhanced, helping us to realize the vision of connecting diffuse scattering to functionally important motions.

Ideas from thermal diffuse scattering theory^35^ support the possibility that sampling of diffuse data might preferentially select for different types of motion. As noted by Peck et al.,^14^ when motions are coupled across unit cell boundaries (as both the present study and their study suggest might be happening in real crystals), the diffuse intensity becomes tied to the Bragg peaks. This effect is closely related to thermal diffuse scattering theory in which the intensity has local maxima at Bragg peak positions and decreases with distance from the peak in a way that is determined by the spectrum of crystal vibrations.^35 14 15 16 2 17^ Motions coupled on long length scales generate intense features that decay sharply moving away from the peaks, and motions coupled on a shorter length scale contribute less intense features that are spread out over a larger region of reciprocal space, extending farther from the peaks. Because the coarse sampling in our study rejects intensity values within ¼ of a Miller index of each Bragg peak, it is dominated by the intensity far from the peaks, enriching the signal due to shorter length-scale correlated motions. In contrast, the finer sampling used by Peck et al.^14^ includes the data corresponding to long length-scale correlated motions, which, despite the localization to fewer grid points near the Bragg peaks, might dominate the fitting due to the higher intensity. In the case of the fine-grained sampling, it is possible that motions on both length scales might be resolved if a LLM with both a short-range and long-range exponential were used.^36^

Although the AMBER and CHARMM simulations yielded similar exponential behavior for the covariance of all Cα atom pairs, a number of differences between the force fields were revealed in our study. (1) The MD simulation pressure after initial solvation was less negative in the case of AMBER (−1439 +/− 39 bar) than CHARMM (−1795 +/− 252 bar). (2) In both Figs. 3 and 4, the covariance in the case of AMBER stays below zero except at the longest distances, and in the case of CHARMM gradually rises to zero after a minimum at around 55 Å. (3) The short-range behavior differs for the AMBER vs CHARMM simulations. For the CHARMM simulations (Fig. 4) the exponential behavior continues to short distances. In contrast, for the AMBER simulations, the covariance in the 3.31 Å bin is negative: −1.77 +/− 1.23 Å^2^. (4) The diffuse intensities predicted from the CHARMM simulation had a somewhat higher correlation with the data than those from the AMBER simulation (Supplementary Fig. S1). At this time the origin of the differences is not clear; however, these differences indicate ways in which crystalline MD simulations, including comparisons to diffuse scattering data, have the potential to distinguish between force fields, and therefore might be used to increase force field accuracy.

There are some caveats to consider in interpreting our results. For all but the inter-protein atom pairs, the covariance increases sharply below 5 Å, deviating from the values predicted by both the LLM and RBM models. Such deviation is not surprising, as short-range interactions are more sensitive to the details of the chemical environment, and include interactions between sequential Cα atoms across the rigid peptide bond. The MD simulations were conducted while constraining the distance between all bonded atoms using LINCS constraints, which further rigidifies the structure, also tending to increase the covariance. Another caveat is that the exponential fit includes a constant term -- the correlation function does not decay to zero as distance increases, perhaps due to the translational alignment of trajectory snapshots.^27^ Note, however, that such a constant offset corresponds to a constant long-range covariance which focuses the diffuse intensity directly beneath the Bragg peak. In this way, the long-range component of the diffuse intensity becomes indistinguishable from the Bragg intensity and does not appear in the measured diffuse intensity. A third caveat is that our analysis was performed using only Cα atom pairs, and therefore does not take into account the influence of non-Cα backbone or side chain motions on the covariance behavior. In particular, it is possible that the signature of RBM might not be as clear for side chain atoms as for Cα atoms, if the backbone is more rigid than the side chains. In future work it will be especially important to analyze simulations in which the bond constraints are relaxed and to overcome the computational difficulties of adding all heavy atoms, including side chain atoms, to the covariance analysis.

Meinhold and Smith^37^ performed an analysis of correlated displacements between atom pairs in MD simulations of crystalline staphylococcal nuclease. Instead of analyzing the covariance matrix, however, they analyzed the dependence of elements of the correlation matrix on the distance between atom pairs. The correlation matrix is computed by renormalizing the covariance matrix, dividing each element by the geometric mean of the values on the diagonal (variances) in the corresponding row and column for each element. Meinhold and Smith found that the correlations of all atom pairs decreased exponentially with a decay length of 11 Å, and that inter-protein atom pairs also decreased exponentially, with a longer decay length (11-18 Å, depending on the simulation). In this respect, the results of our analyses are similar. However, in contrast to the near linear behavior we found for the covariance of intra-protein atom pairs, they found that the intra-protein correlations showed an exponential decay behavior. Our analysis therefore appears to be inconsistent with theirs with respect to the intra-protein atom pairs. We note that there are several differences between our analyses: on the one hand, Meinhold and Smith^37^ analyzed the correlation matrix, used a single periodic unit cell, a NPT ensemble, and 10 ns duration simulations; on the other hand, we analyzed the covariance matrix, used a 2×2×2 periodic supercell, a NVT ensemble, and 600 ns duration simulations. It is possible that the discrepancy with respect to the intra-atom pairs is due to one of these differences.

As mentioned above, the correlation of the anisotropic intensity calculated from the LLM model with the data is 0.73, which is higher than the maximum value of 0.63 for the MD simulation. The higher correlation of the LLM suggests that it provides a globally more accurate description of the anisotropic intensity. Although the MD model is lacking in this respect, it still has advantages over the LLM model. For one, the MD simulation is still the only model that is capable of reproducing the total intensity – isotropic and anisotropic – and is therefore the most accurate overall. Moreover, the MD model is, in some sense, a “model-free” model, in that there are no free parameters to fit – the model simply depends on the choice of force-field, and the assumption that the system behaves classically. Indeed, the LLM model accuracy is substantially increased due to the ability to refine the free parameters against the experimental data. However accurate the LLM model may be, it produces a very limited description of the dynamics that does not contain any mechanistic information, whereas the MD model can provide us with dynamic structural information that yields functional and biological insight (modulo any inaccuracies inherent to the forcefield, or due to inadequate sampling and/or simulation length).

Taken together, our results show that MD simulations of a crystalline protein exhibit LLM behavior. The inter-protein atom pairs exhibit LLM behavior and within the protein the motions exhibit both LLM and RBM behaviors. Due to the large number of inter-protein vs intra-protein atom pairs, the overall behavior appears LLM-like. These findings provide support and context for previous results which showed that LLM models of protein diffuse scattering improve after the inclusion of interactions across protein boundaries. They also provide clues about why LLM model fits using coarsely sampled diffuse data might yield smaller correlation length scales than using finely sampled data. Finally, our results suggest that the modeling of finely sampled diffuse scattering data might be improved by consideration of both small-scale and large-scale collective motions.

## Methods

### Molecular-dynamics simulations

The all atom structure for staphylococcal nuclease was pulled from the Protein Data Bank (wwPDB: 4WOR^38^ with unit cell *a* = *b* = 48.5 Å, *c* = 63.43 Å, α = β = *γ* = 90 degrees, and space group P41). This structure is missing the first five residues at the N-terminus and the last eight residues at the C-terminus. In Ref. 28 the missing N- and C-terminal atoms were reintroduced and modelled based on extension of secondary structure – the same starting structure is used in this work.

Once fully modelled, the asymmetric unit was propagated to a unit cell and then to a 2×2×2 supercell using the *UnitCell* and *PropPDB* methods from AmberTools18.^39^ The coordinates of the bound ligand, thymidine-3′-5′-bisphosphate (pdTp), were extracted from the PDB file, saved as a mol2 file (using UCSF *Chimera*^30^), and parameterized using the SwissParam Server (*swissparam.ch*^40^). Two different systems were created: one in which the protein residues were parameterized with the AMBER 14SB force field^41^ and another in which they were parameterized with the CHARMM 27 force field^42^; both were parameterized using GROMACS^43^ *pdb2gmx* (residue names were set manually and hydrogens present in the initial PDB file were ignored with flag *-ignh*, which automatically assigns protonation states for residues at pH 7). These fully parameterized systems were then solvated with TIP3P waters^44^ using GROMACS *gmx solvate*. The full systems were neutralized with chloride ions (*gmx genion)*. Once solvated, these systems were minimized using the steepest descent algorithm.

Simulations were performed using a constant particle number, volume, and temperature (NVT) ensemble, at a temperature of 298K. After an initial round of NVT equilibration to check the pressure of the system, the number of water molecules was adjusted to achieve near-atmospheric pressure. This was achieved by iterative rounds of solvation and NVT equilibration. For the CHARMM force field simulation, 100 ps equilibration durations were used, as in previous studies of the same system.^25, 28^ For the AMBER force field simulation, 5 ns equilibration durations were used. After the last round of equilibration, 17,557 waters were added in the AMBER simulation, and 17,138 waters were added in the CHARMM simulation.

The crystallographic protein heavy atoms (i.e. non-terminal heavy atoms) were restrained to the minimized crystal structure during all rounds of equilibration and the initial 100ns restrained production simulation (restraint force constant k=1000 kJ mol^−1^ nm^−2^). Restraints were then released and production simulation was carried out for 600ns. All rounds of equilibration and production were carried out using the leap-frog algorithm (*integrator=md*); neighbor searching was carried out using the Verlet cutoff-scheme^45^ with an update frequency of 10 frames (*niter=10*), and a cutoff distance for the short range neighbor list of 1.5 nm (*rlist=1.5*); all bonds were constrained with the LINCS algorithm (*constraints=all-bonds; constraint-algorithm=LINCS).*^46^

### Covariance matrix of atom displacements

After releasing restraints, the simulations require on the order of 100ns for the RMSD of the protein Cα coordinates to plateau. To ensure that the system was fully equilibrated after the release of restraints, analysis began at 200ns in to unrestrained production. Cα trajectory subsets were extracted from the full unrestrained production trajectories from 200-600ns in both simulations. To do this, the first frame (200ns into unrestrained production) was extracted, and the Cα coordinates were isolated using *gmx editconf* (with flag *-pbc* to ensure molecules stay whole). Then, a 400ns Cα trajectory was created using *gmx trjconv*, and subsequently translationally fit to the starting structure using the Cα starting frame as reference (*gmx trjconv … -s c_alpha_supercell_pbc.gro … -fit translation)*. The Cα trajectory covariance information was calculated using *gmx covar* (once again, using the Cα structure from the first frame as reference).

With 32 proteins, each containing 149 Cα atoms, and a 3 by 3 covariance submatrix for each pair of Cα atoms, the full covariance matrix computed by *gmx covar* is a 14,304 by 14,304 block matrix. The diagonal elements of this matrix correspond to the mean squared deviation (MSD) for each atom, in each direction; these diagonal elements were ignored in subsequent analysis. After computing the trace of each atoms-pair’s 3 by 3 submatrix, the covariance matrix is 4,768 by 4,768 (Supplementary Fig. S2). These pairwise Cα covariances were sorted by their distances apart, using a matrix of pairwise Cα distances (Supplementary Fig. S2) computed using *MDTraj*^47^ and the average coordinates reported by *gmx covar* for each Cα trajectory subset. Covariance matrix data was processed, analyzed, and plotted using python’s *numpy, scipy, and matplotlib.* The covariance as a function of distance data was fit to an exponential decay model using *gnuplot*, and errors in the parameters reported in the Results section are the asymptotic standard errors reported by *gnuplot*.

### Rigid-body motions model

To investigate the source of the non-exponential decrease in covariance as a function of distance for residue pairs within proteins, we created a Python script to simulate RBM (*RigidBodyMotions.py*). The script creates hypothetical trajectories consisting of a rigid protein randomly rotated and translated by amounts on par with the magnitude of motion observed in our MD simulations. We compared the covariance behavior computed from these "trajectories" with that observed in the crystalline MD simulations. This allowed us to determine the degree to which a rigid-body rotation and translation model can explain the MD covariance behavior.

The Python script generates covariance data as follows:

1. The structure of a single protein is pulled from the crystalline MD starting structure, and centered on the origin (using *mdtraj.Trajectory().center_coordinates()*); terminal atoms missing from the PDB structure are disregarded, as they cannot reasonably be considered rigid.
2. Rotations are generated by sampling three Euler angles, each from a normal distribution with mean zero and a specified standard deviation; similarly, translations are generated by sampling a three-dimensional vector from a normal distribution with mean at the origin and a specified standard deviation, the same for all directions.
3. A new “frame” is created by first rotationally moving and then translationally moving the starting structure; for rotational moves, a rotation matrix is generated from the Euler angles, and the matrix vector product of the rotation matrix and the coordinates of each atom is performed (with *numpy.dot()*); for translations, the random three-dimensional translation vector is added to each atom’s coordinates.
4. A “trajectory” is built up frame by frame, and the covariance is calculated as a function of the distance between atoms, as described above for simulation trajectories.

In this study we used 5000 frames for our analysis. The covariance data produced was compared with the covariance data from equivalent atoms in the simulation (non-terminal Cα atom pairs within proteins). Reasonable parameters for the Euler angle and translational distributions were arrived at by manual adjustment of the standard deviations, seeking the best visual overlap between the model and the data.

### Diffuse scattering

Diffuse scattering data for staphylococcal nuclease were obtained from past experiments^38^ and were processed as described in Ref. 28. In addition to studying the liquid-like motions behavior of the atom displacement covariance matrix, the MD trajectories themselves can be processed directly to predict the diffuse scattering. The diffuse scattering is the variance of the structure factor of independent repeating units in the crystal, according to Guinier’s equation,^48^ and previous studies have predicted the diffuse scattering from protein crystal MD trajectories by computing the structure factor, frame-by-frame.^25, 27, 28, 37, 49^ This is done as in Ref. 28 using the script *get_diffuse_from_md.py*, which takes in a trajectory and outputs *.mtz* (byte stream) and *.dat* (ascii) reflection data; then, reflected data is processed with the *Lunus* diffuse scattering data processing software suite (https://github.com/mewall/lunus). The same script and processing software were used in this work.

Diffuse scattering simulations were calculated from the last 400ns of the trajectories. As in Ref. 28, the intensities were computed on a 3D grid sampling reciprocal space twice as finely as the Bragg lattice. The trajectory was produced in 100ns chunks, and the diffuse scattering was calculated from these 100ns chunks independently, and then accumulated either (a) *coherently*, by accumulating the complex structure factors (flag *-merge=True*) or (b) *incoherently*, by averaging the intensities themselves from each 100ns chunk using *Lunus* methods *sumlt* and *mulsclt*. The model diffuse scattering was converted to a lattice file using *hkl2lat*, with the experimental lattice file as a template, then symmetrized and culled using *symlt* and *culllt*, and the anisotropic component of the diffuse scattering was separated using *anisolt*. Pearson correlations between the models and the data were computed using *corrlt*.

Simulated diffraction images in Fig. 2 and Supplementary Fig. S4 were computed from models or data in a similar way to that in Ref. 28. To simulate diffraction using a 3D grid model or data, the minimum value was computed within shells and was subtracted from each 3D grid point, using interpolation (*subminlt*). The diffraction pattern corresponding to a specified crystal orientation was simulated from the 3D grid using the *simulate_diffraction_image.py* Python script distributed with *Lunus*.

For the refinement of the LLM, we processed the data using a recent version of *Lunus* (https://github.com/mewall/lunus). As in the original staphylococcal nuclease study ^38^, the data were coarsely sampled using one point per Miller index. The data were processed to a resolution limit of 1.6 Å. Intensity values within ¼ of a Miller index of each Bragg peak were excluded from the processing. The 3D data were symmetrized using the P4 Laue symmetry, and the isotropic component was removed as described in Ref. 28. The structure factors from the 4WOR crystal structure were used to refine a LLM model (*python refine_llm.py symop=-3 model=llm bfacs=zero*).

## Abbreviations

MD: molecular dynamics
LLM: liquid-like motions
RBM: rigid-body motions
pdTp: thymidine-3′-5′-bisphosphate

## Acknowledgments

This work was supported by the University of California Laboratory Fees Research Program. MEW is also supported by the Exascale Computing Project (17-SC-20-SC), a collaborative effort of the U.S. Department of Energy Office of Science and the National Nuclear Security Administration. JSF is also supported by NSF STC-1231306. The simulations were performed using Institutional Computing machines at Los Alamos National Laboratory, supported by the U.S. Department of Energy under Contract 89233218CNA000001.

## Supplementary Information

**Figure S1.**
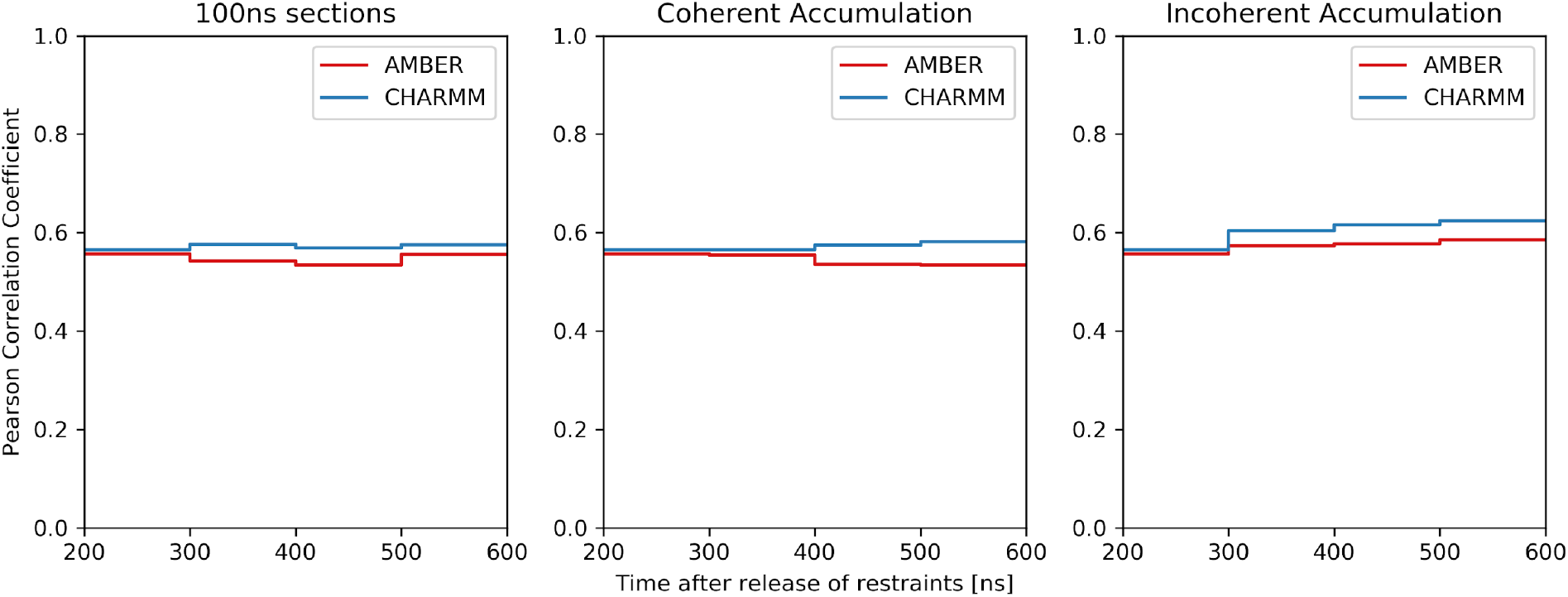
Agreement between 3D diffuse scattering data and simulations computed using sections of the MD trajectory. Values of the Pearson correlation coefficient are computed as a figure of merit. AMBER values are in red, CHARMM in blue. *Left*. Agreement computed using individual 100 ns sections after the 200 ns time point (200-300, 300-400, 400-500, and 500-600 ns). Values do not systematically increase or decrease with the simulation time. *Center*. Simulations computed using trajectories accumulated coherently after the 200 ns time point (see Methods for definition of coherent accumulation): (200-300, 200-400, 200-500, and 200-600 ns). The AMBER values decrease and the CHARMM values increase with increasing time. *Right.* Simulations computed using trajectories accumulated incoherently across 100 ns sections after the 200 ns time point (see Methods for definition of incoherent accumulation). Both the AMBER and CHARMM values increase with increasing time. Incoherent accumulation results in the highest correlation with the diffuse scattering data, and the CHARMM correlation is higher than the AMBER correlation in all cases.

**Figure S2.**
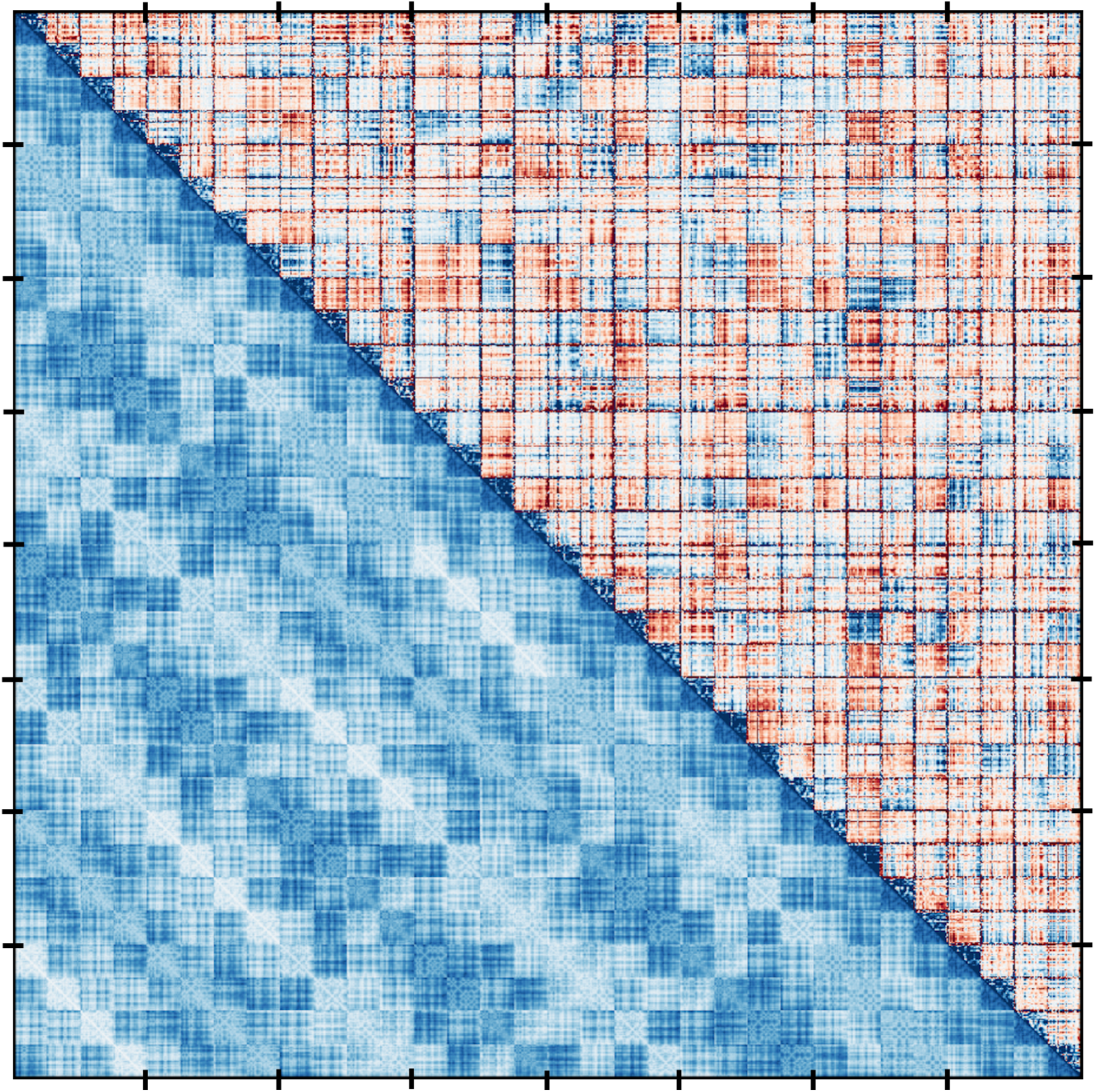
Visualization of values from the Cα atom displacement covariance matrix (*upper right triangle*) and the Cα atom distance matrix (*lower left triangle*). Rows and columns correspond to Cα atoms (the image shows only a sampling of the values from the full matrix and merely illustrates the overall structure of the data, as the resolution is insufficient to show values for all atom pairs). Tics indicate breaks between each of eight unit cells. The finer squares within each unit cell correspond to four copies of the proteins, each of which has 149 Cα atoms. The covariance values in the upper right triangle range from −0.2 (*dark red*) to 0 (*white*) to + 0.2 (*dark blue*), with values above or below in absolute value visualized as the same dark blue/red color. The distance values in the lower left triangle range from 0 (*dark blue*) to 92 Å (*white*). The blocks along the diagonal correspond to intra-protein atom pairs; for these blocks, the covariance tends to be positive (*blue*) and the distances are short (also *blue*). All other blocks correspond to inter-protein atom pairs; for these blocks, there are more negative covariance values than for the intra-protein blocks.

**Figure S3.**
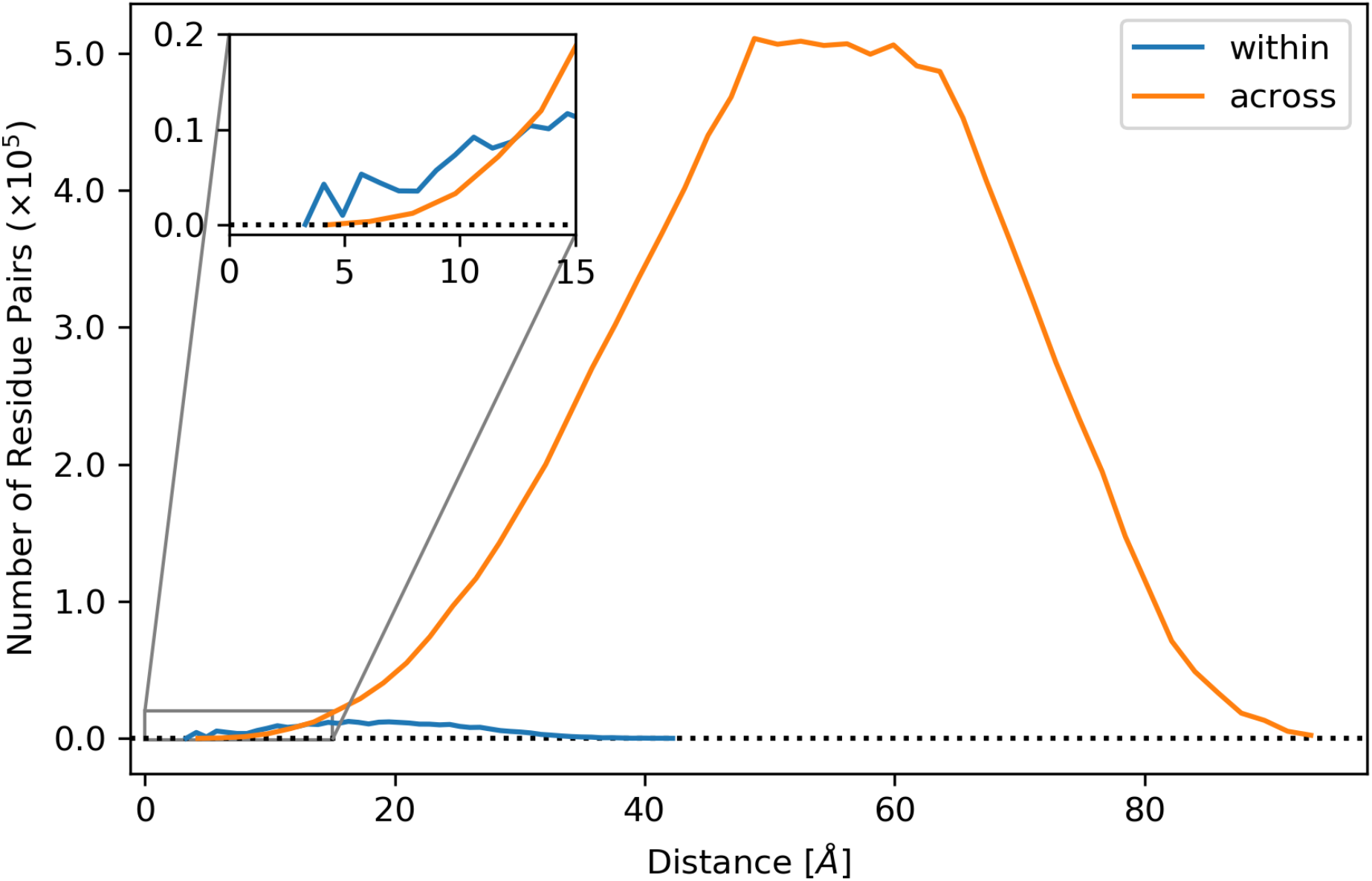
Number of Cα atom pairs as a function of distance in the full supercell. Intra-protein (within) atom pairs are shown in blue; inter-protein (across) atom pairs are shown in orange. The number of pairs across proteins outnumber the residue pairs within proteins for all distances greater than 12 Å (*inset*).

**Figure S4.**
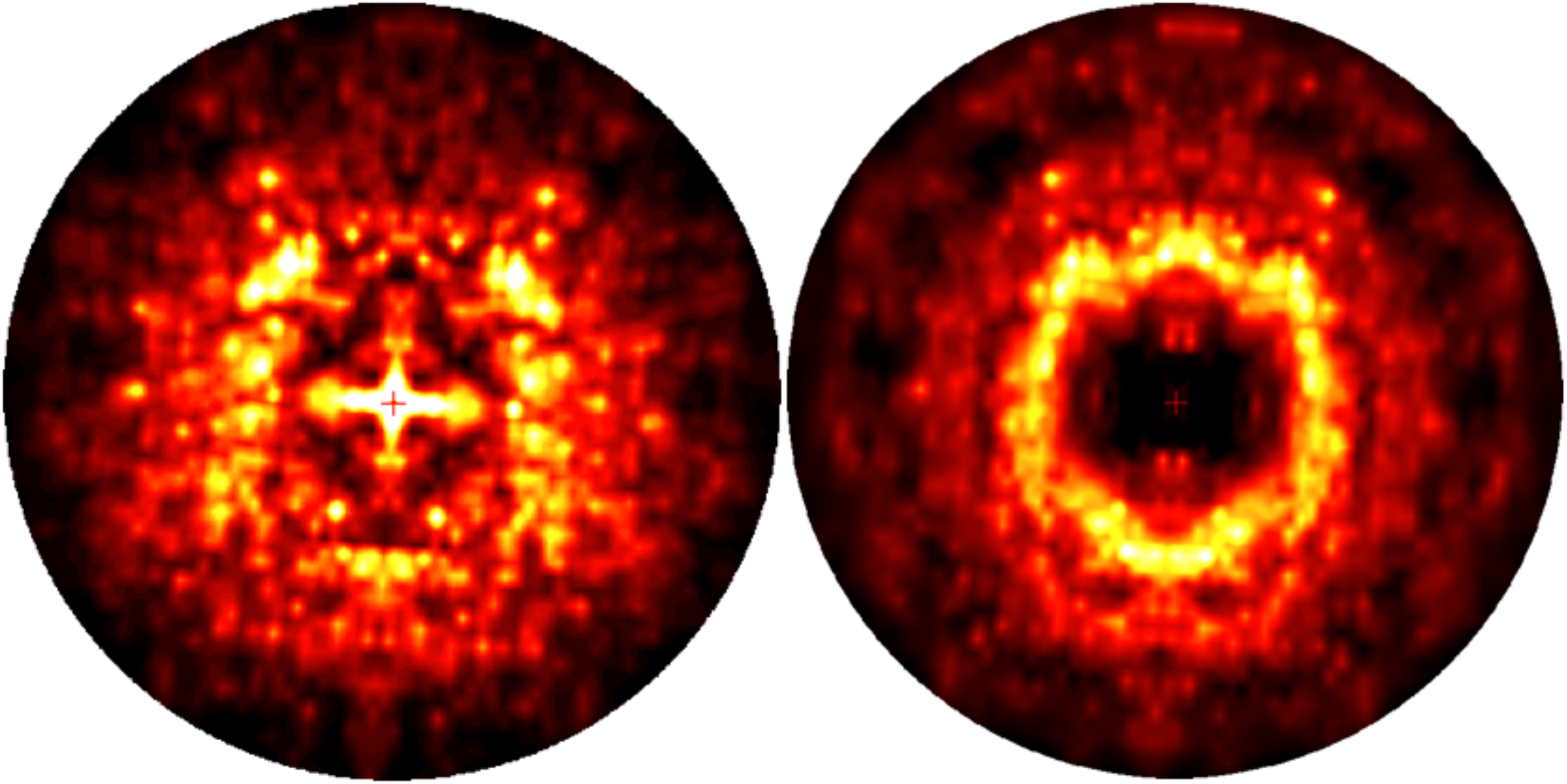
Simulated diffraction images from LLM model (*left*) and 3D experimental diffuse data (*right*). Both the data and model were generated using coarse sampling of one measurement per Miller index. Outside of the region at low resolution near the origin (which, despite the small contribution of this region in determining the quantitative agreement, can dominate the visual comparison due to its central position), similar features are identifiable between the two images. The images were displayed using the heat map mode of ADXV.^31^

